# Arterial blood contrast (ABC) enabled by magnetization transfer (MT): a novel MRI technique for enhancing the measurement of brain activation changes

**DOI:** 10.1101/2020.05.20.106666

**Authors:** Jenni Schulz, Zahra Fazal, Riccardo Metere, José P. Marques, David G. Norris

## Abstract

Functional brain imaging in humans is almost exclusively performed using blood oxygenation level dependent (BOLD) contrast. This typically requires a period of tens of milliseconds after excitation of the spin system to achieve maximum contrast, leading to inefficient use of acquisition time, reduced image quality, and inhomogeneous sensitivity throughout the cortex. We utilise magnetisation transfer to suppress the signal differentially from grey matter relative to blood so that the local increase in blood volume associated with brain activation (mainly occurring in the arterioles and capillaries) will increase the measured signal. Arterial blood contrast (ABC) is additive to the residual BOLD effect, but will have its maximum value at the time of excitation. We measured brain activation using combined ABC and residual BOLD contrast at different times post-excitation and compared this to BOLD data acquired under otherwise identical conditions. We conclude that using ABC and measuring shortly after excitation gives comparable sensitivity to standard BOLD but will provide greater efficiency, spatial specificity, improved image quality, and lower inter-subject variability. ABC offers new perspectives for performing functional MRI.

## Introduction

Brain activation studies conducted using functional magnetic resonance imaging (fMRI) have made an enormous contribution to our understanding of the biological underpinnings of human cognition and behaviour, as well as having a broader societal impact, ranging from pre-operative planning, diagnosis, to improved understanding of mental illnesses and neurological disorders.^1^

fMRI typically uses Blood Oxygenation Level Dependent (BOLD) contrast whereby paramagnetic deoxyhemoglobin acts as an endogenous contrast agent.^2^ Upon brain activation, local changes in blood flow (CBF), -volume (CBV), and metabolic rate lead to a reduction in deoxyhemoglobin concentration and a corresponding increase in the transverse relaxation time. The resultant signal changes are measured most commonly with a gradient-echo echo planar imaging (GE-EPI) sequence. For optimum sensitivity, the echo time (TE) should match the transverse relaxation time of the tissue, typically some tens of milliseconds. ^3^ This technique has the following disadvantages: First, the relatively long TE limits the efficiency, reduces the signal intensity by a factor of about three, and makes the method prone to artefacts, particularly signal dropout in regions of poor static main field homogeneity. Second, reliance on changes in deoxyhaemoglobin concentration in the capillaries and post-capillary vessels has the consequence that the measured activation will always be downstream of the underlying neuronal activity. Third, the sensitivity will vary through the brain with the local value of the transverse relaxation time.

Recently there has been increasing interest in measuring brain activation by using techniques based on changes in CBV,^4-13^ which is known to be better localised to the source of neuronal activity than BOLD.^14, 15^ One approach to sensitise the MRI signal to changes in blood volume is to use magnetisation transfer to differentially attenuate the signal from tissue, while that from blood is largely unaffected.^16, 17^ An increase in blood volume, such as that occurring as a result of brain activation, will lead to an increase in signal intensity.^18-20^ Early applications of MT, in conjunction with brain activation studies, explored the effect of MT on BOLD-contrast,^21, 22^ or exploited MT effects to measure blood volume.^18, 20^ These experiments used lengthy off-resonance pulses of several seconds duration and TEs similar to those used in standard BOLD imaging.

Here, we diverge from previous approaches by using on-resonance pulsed MT^23^ and minimising TE. Pulsed MT efficiently saturates tissue signal,^23^ increasing the sensitivity to changes in blood volume, which will be highest at TE=0. The combined arterial and capillary contribution to the blood volume changes has been estimated to be 80-90%.^24, 25^ Water exchange between the tissue and capillaries will also diminish the venous contribution when MT partially saturates the tissue signal.^19^ This will reduce the relative contribution of the venous compartment to the signal. We can thus expect at least 80% of the MT-weighted signal variation at TE=0 to come from arterioles and capillaries. At rest, it has been shown that most oxygen exchange occurs in the smallest arterioles, and in the first branches of the capillary bed.^26^ Upon activation, the increases in arterial and capillary blood volume will lead to increased flow and perfusion. If we suppress the tissue signal using MT, then we can expect an increase in the total signal to occur upon activation because of increased blood volume, but also because of an increase in the water exchange between tissue and capillaries. For this reason, we have elected to term the additional functional contrast generated by suppression of grey matter signal ‘arterial blood contrast’ (ABC).

One fortunate characteristic of ABC contrast is that it is additive to the BOLD response, as an increase in brain activity will lead to an increase in both signals. With ABC, the signal increase can be ascribed to an increase in the magnetisation within a voxel. Whereas with BOLD, the signal increases are caused by an increase in the transverse relaxation time. To estimate a pure ABC signal change, it would be necessary to perform an experiment at zero TE, which is technically challenging, especially within the constraints of a realistic fMRI experiment. We can expect, however, that the introduction of ABC will modify the activation recorded at short TEs. In this work, we used a multi-echo gradient-echo echo-planar imaging (EPI) pulse sequence with and without MT applied to acquire data over a range of TE-values. The two experiments are referred to as MT-on and MT-off, with the TE for data acquired at a given TE denoted in subscript, *e.g.*, MT-on_TE=6.9_ for the first echo of the MT-on acquisition. By keeping all other parameters constant between the two datasets, we explored whether the application of MT improves the sensitivity to activation at short TE, relative to the BOLD response, which we use as a benchmark. Specifically, the images from the MT-on experiment can be expected to have an ABC that will decay exponentially with TE, and a BOLD contribution that will increase with TE. The MT-off experiment will display the classic BOLD signal dependence on TE,^3^ with full BOLD sensitivity.

## Results

Data were obtained from 16 subjects using a visual stimulus paradigm (4 Hz flickering chequerboard), pre-processed using standard fMRI analysis tools, and a first level statistical analysis was performed (*c.f.* Methods). Group level activation maps were obtained for both fixed- and mixed-effect statistics,^27^ significance levels were set following recommended procedures.^28^ Figure 1 shows the results of the fixed effects analysis. In figure 1a, we show this in the form of activation maps from four representative slices. Significant activation is obtained at all echo times for both contrasts as would be expected. At each TE the activation appears stronger and more widespread for MT-on than at the corresponding TE for MT-off. To elucidate further these activation patterns, we defined the activated region of interest (ROI) as the union of the MT-off_TE=28_ and MT-on_TE=6.9_ group-level activation maps. Figure 1b shows the mean z-scores of activated voxels within this region as a measure of functional sensitivity, as well as the total number of significantly activated voxels (both thresholded at z>3.1). Both contrasts show monotonically increasing z-scores as a function of TE. The highest z-scores are recorded for MT-on_TE=18_ and MT-on_TE=28_. The number of activated voxels is higher for all MT-on experiments than for MT-off at the same TE.

**Figure 1.**
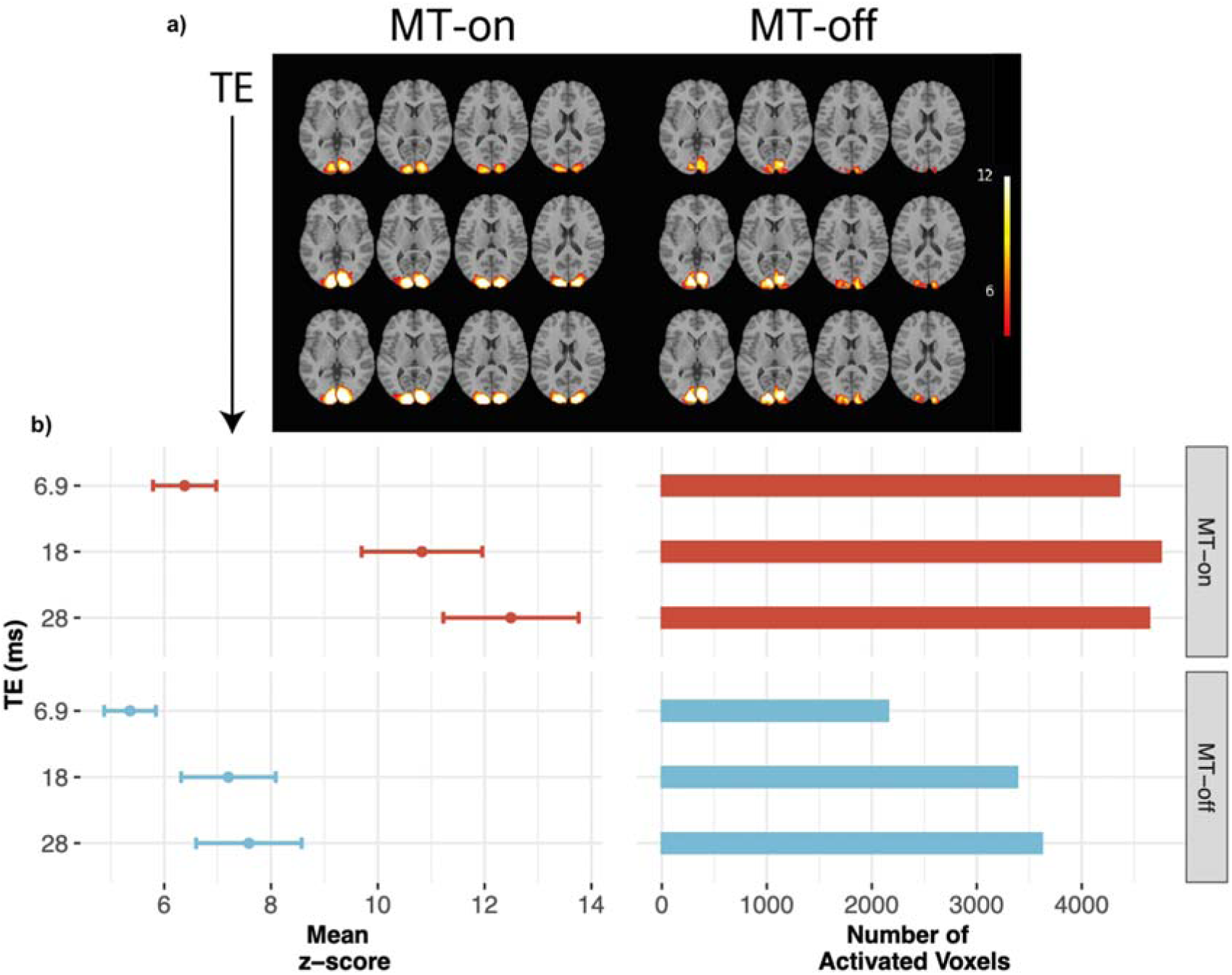
a) Fixed effects group-level activation maps within the visual cortex covering four slices, for all three TE-values (6.9/18/28ms). b) Mean z-scores and the number of activated voxels within the ROI defined by the union of MT-on_TE=6.9_ and MT-off_TE=28_ activation map at group-level (Z>3.1). The values are shown only for pixels with significant activation at Z>3.1

A mixed effects analysis shows the response to the task without the inter-subject variation, which is modelled separately. The *z*-scores are correspondingly reduced compared to fixed-effects, as shown in figure 2. For MT-off, no significant activation is recorded at the two shortest echo times. All MT-on data showed significant activation at all TEs but with less activated voxels than for MT-off. The difference between the *z*-score at the shortest TE and those for the two longer TEs is also reduced compared to the fixed effects analysis.

**Figure 2.**
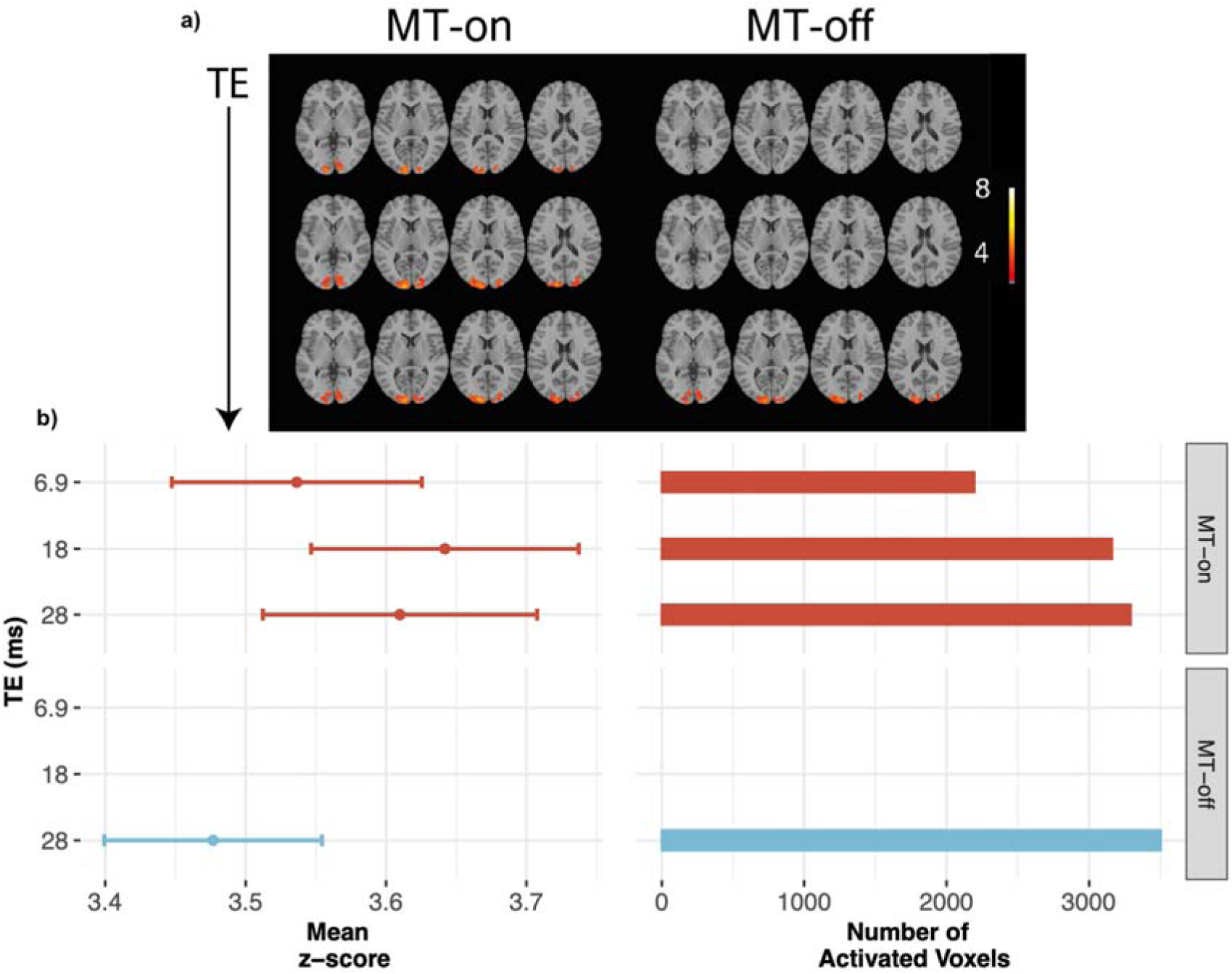
a) Mixed effects group-level activation maps within the visual cortex covering the same four slices as figure 1, for all three TE-values (6.9/18/28ms). b) Mean z-scores and the number of activated voxels are calculated within the ROI defined by the union of MT-on_TE=6.9_ and MT-off_TE=28_ activation map at group-level (Z>3.1). Values are shown only for pixels with significant activation at Z>3.1.

## Discussion

These results provide convincing evidence for an underlying contrast mechanism that differs from BOLD. This argument is based on how the statistical maps vary as a function of TE, and the difference between the results obtained with fixed- and mixed-effects analyses. For BOLD, the z-scores increase with TE as expected for a fixed effects analysis, whereas for mixed effects only at TE=28 ms, is significant activation detected. For MT-on, the pattern is markedly different. Although significant activation is recorded at all TEs, in both fixed and mixed effects analyses the relative strength of activation at shorter TEs increases for mixed effects, in marked contrast to BOLD. This behaviour is consistent with the presence of a qualitatively different contrast. The most plausible mechanism for the additional signal change is that it is driven by changes in CBV in the arterioles and capillaries, *i.e.* the ABC mechanism proposed in the Introduction. This would also be expected to give more consistent activation patterns between subjects than BOLD, where additional variance can be induced by the signal contribution downstream from the region of activation, thus explaining the improved performance in the mixed effects analysis.

The MT pulses are not slice selective, and in the present work are transmitted via the body coil, and hence inflow can be excluded as the source of the ABC-contrast. It is highly likely that the application of MT will change the physiological noise characteristics. However, a change in noise level is an implausible explanation of the qualitative changes in behaviour that we observe.

Given the strong tissue signal reduction caused by MT, it is at first sight seem surprising that activation increases with TE for MT-on. This is however, a consequence of the ABC mechanism: the increase in CBV will give rise to a concomitant decrease in the tissue volume. This decrease is what is utilised in the VASO technique to examine changes in CBV.^4^ Activation reduces the tissue magnetisation but increases the T2*. The tissue signal change upon activation will be maximally negative at TE=0 and increase in value with increasing TE. Hence reducing this negative signal will increase the functionally induced signal change at short TEs. Although the ABC contrast at TE=0 will have a small venous contribution, the residual BOLD signal recorded at shorter TE-values will have a higher relative intra-vascular contribution. These effects are explored quantitatively in supplementary information.

There is a further effect, which may contribute beneficially to the activation recorded during MT-on. It has previously been discussed that the exchange of water over the capillary wall could reduce the post-capillary signal. ^19^ The corollary is that the unsaturated magnetisation entering the tissue will enhance its signal. The exchanging magnetisation will of course equilibrate with its surroundings on the time scale of T1, but the brain is a highly perfused organ, and at rest, about 1% of the water is exchanged between tissue and blood per second, which will increase to about 1.6% upon activation.^29, 30^ The consequence of this exchange would be to reduce the intravascular signal while increasing the tissue signal in the vicinity of the capillaries. This would additionally improve the spatial specificity of signal changes measured upon activation. This mechanism is similar for that recently proposed for the combined measurement of blood volume and perfusion changes obtained by suppressing the signal of inflowing blood.^31^

Our approach relies on two hitherto unexplored aspects of MT as applied to functional neuroimaging: pulsed MT and minimal TE. We have implemented a pulsed MT-block prior to each excitation pulse, although a larger interval of about 100 ms may be optimal. ^32^ If we would implement our MT-block using this interval then scanning time would increase by only 6%, and there is considerable scope for reducing this further by using shorter RF pulses to induce MT.^33, 34^ In addition to the efficient MT contrast generation, the use of a short TE increases both sensitivity and efficiency. It also allows us to reduce or eliminate the signal dropouts that manifest at long TE in regions of poor static main field homogeneity, and means that fMRI no longer requires long TE experiments, opening new possibilities for data acquisition, such as spiral trajectories^35-37^ and short TE implementations of both 2D-^38, 39^ and 3D-EPI.^40, 41^ Although we chose the visual cortex for our initial implementation because of the strong activation that can be elicited in this region, there is no *a priori* reason to believe that our results are not applicable to the whole brain. ABC will have improved spatial localisation as it is mainly based on arterial CBV changes. Thus, high-resolution fMRI studies will benefit from a high ABC contribution to the functional contrast. Variations in MT efficiency through the brain (*c.f.* Figure S2) are far lower than the variation in T2*. Hence, by appropriate application of MT, all brain regions should become more equally accessible. The marked improvement in mixed effects statistics that we observe will be particularly beneficial for larger group and population studies. As ABC is complementary to BOLD and straightforward to implement, we expect it to achieve widespread use in the near future.

## Methods

### MT On-resonance pulse

To suppress the tissue signal, a net 0° on-resonance MT preparation block^23^ was implemented into the multi-echo EPI sequence. The MT preparation block consists of two non-selective binomial RF pulses of opposite phases and magnitudes in the proportions ±1/ ∓2/ ±1 (6 ms). It is followed by a pseudo-random gradient spoiler scheme (3 ms) in all three directions (see figure 3a). The application of two binomial pulses back to back with inverted phase is aimed at reducing on-resonance signal attenuation, which could be related to transmit amplitude errors or large off-resonance effects due to static background field inhomogeneity.^42^ Piloting showed that multiple repetitions of the dual-binomial MT pulse did not show any further improvement in MT image quality. The pseudo-random gradient was designed to avoid coherence pathways associated with the signal excited by the MT pulse (for example, fat signal). To achieve a constant steady state of continuous tissue suppression, the MT-block was played out before each slice excitation. The MT-flip angle pulse amplitude and pulse duration were optimized in a preliminary experiment to attenuate maximally grey matter signal while restricting the attenuation of CSF to about 10%. We used the CSF signal attenuation in the ventricles, as a proxy for the arterial blood signal attenuation. Experiments were typically performed at about 60% of the permitted specific absorption rate (SAR), of which approximately 95% could be attributed to the MT-pulses.

**Figure 3:**
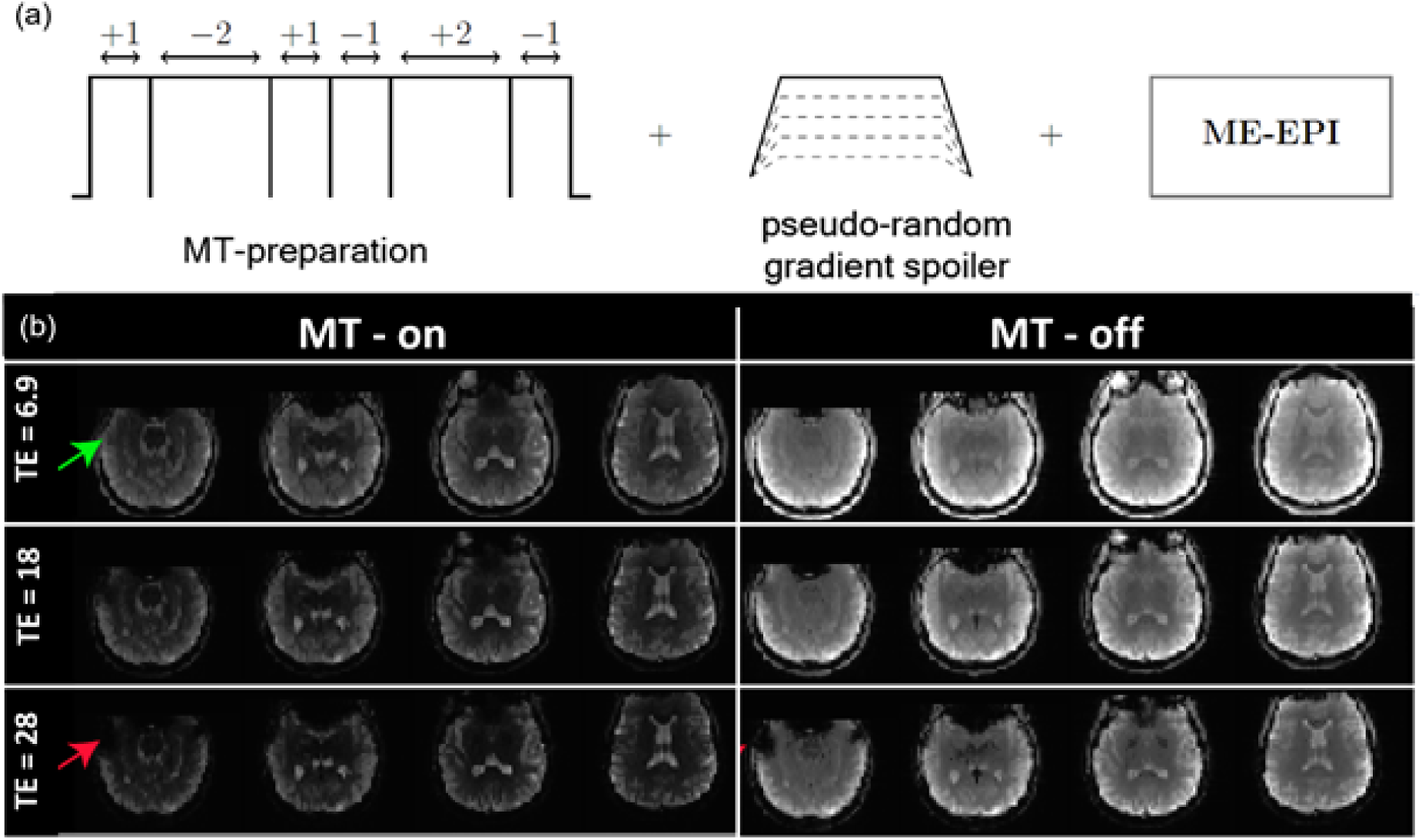
(a) Sequence design with the MT preparation module (6ms) and the pseudo-random gradient spoiler scheme (3ms) played out before every excitation of the ME-EPI sequence. (b) The image quality for MT-on and MT-off at different echoes. The green arrow indicates reduced dropout for MT-on_TE=6.9_ as compared to MT-off_TE=28_. Red arrows indicate dropout regions for MT-on/off_TE=28_

### Scanning protocols

The choice of protocols was determined based on preliminary experiments. ^43, 44^ The MT-pulse angle was adjusted empirically to maximally attenuate grey matter signal while restricting the attenuation of CSF to about 10%. Data were acquired on a Siemens MAGNETOM Prisma (3T) MRI scanner (Erlangen, Germany) with a 32-channel head coil. The acquisition was based on a multi-echo gradient-echo 2D-EPI with MT-on and MT-off conditions. In the MT-on condition, the MT-block was implemented as explained above, whereas, in the MT-off condition, the RF-pulse of the MT-block was turned off while the timing conditions remained the same. The MT flip angle for the MT-on acquisition was ±77°/∓154°/±77°. For both conditions, we acquired ME data at echo times TE=6.9/18/28ms. The first echo time was chosen to be as short as possible to maximise ABC in the MT-on condition. The upper limit was determined to ensure that a strong standard BOLD signal would be recorded for the MT-off experiment, but that TR would not exceed 2s.

The acquisition protocol for the ME GE-EPI had the following parameters: in-plane resolution 3 mm isotropic and 38 slices without a gap for a coverage of 12 cm, Acquisition matrix 80×80, and fat saturation performed before each RF excitation. The flip angle for RF excitation was based on the Ernst angle of grey-matter (50°). The in-plane acceleration factor was three, with a partial Fourier factor of 6/8 and 2718 Hz/pixel acquisition bandwidth.

Anatomical scans were acquired for image registration using a sagittal 1 mm isotropic MP-RAGE with: TR of 2300 ms, TI of 900 ms, TE of 3 ms, FA 9°, turbo factor 16 and an in-plane acceleration factor of 2. The total acquisition time was 5:20min. All imaging sequences were automatically aligned using an auto-align localizer sequence.

### Data Acquisition

The data quality using MT-on acquisition is shown in figure 3b, with uniform suppression of grey matter signal in the regions of activation (figure S2). We acquired ME-GE EPI data from 16 subjects (10M/6F 26 ± 6 yrs) with this number based on power analysis for a mixed-effects analysis at the group-level.^45^ Pilot-data from two preliminary studies were used for the power calculation (three subjects (2M/1F 28 ± 4 yrs) and 7 subjects (3M/4F 25 ±4 yrs)).^43, 44^ The power analysis was implemented in the fMRIpower software package (http://fmripower.org). This method estimates power for detecting significant activation within specific regions of interest, with the assumption that the planned studies will have the same number of runs per subject, runs of the same length, similar scanner noise characteristics, and data analysis with a comparable model. All power calculations were based on an atlas^46^ and used a *p-*value threshold of 0.005 for a one-sided hypothesis test. With 16 subjects, we predicted at least 80% power to detect an effect size of 0.7 in the Inferior Occipital Gyrus (IOG) with MT-on at TE of 6.9 ms and MT-off at TE=28 ms using mixed-effects analysis at the group-level.

The participants were shown a hemifield (R/L) checkerboard flickering at 4Hz randomly distributed across trials with a block design of [10s on, 26-32 off ISI]. During the entire task, the participants were asked to focus on the grey fixation cross in the middle of the screen and press the button box with the R/L index finger whenever the fixation cross changed colour using the immediately previous stimulus hemifield as a reference for R or L. The total duration of the task was 10 minutes. Stimuli were presented, and button presses were recorded using MATLAB 2018a, (The MathWorks, Inc., Natick, Massachusetts, United States). Before performing the task in the scanner, participants practiced on a desktop computer located next to the scanning console to guarantee that the procedure was understood. Each subject performed two runs of ten minutes during a session with one MT-on and the other MT-off. This is because the lengthy duration required for the build-up of steady-state and decay of the MT effect makes it difficult to alternate in a single run. The order in which MT-on/off was applied was counterbalanced across subjects.

### Functional Processing

Before data pre-processing, DICOMs were converted to NIfTI’s using dcm2niix.^47^ For each BOLD run, the following pre-processing was performed using FSL. First, head-motion parameters were estimated using a reference volume for the BOLD data (transformation matrices, and six corresponding rotation and translation parameters) using MCFLIRT.^48^ The BOLD data were then co-registered to the T1-weighted reference using FLIRT^49^ with the boundary-based registration cost-function.^50^ Co-registration was configured with nine degrees of freedom to account for distortions remaining in the BOLD reference. The BOLD data were resampled to MNI152NLin2009cAsym to generate pre-processed BOLD data in standard space.^51, 52^ These were smoothed with a 6mm kernel and high pass filtered with a cut-off frequency of 1/100s. For MT-on data, the first three volumes (∼6 sec, including all echoes) were removed to ensure a steady-state for MT contrast. The hemodynamic response was evaluated for MT-on and MT-off data, by computing, the hemodynamic responses within the right and left hemifields using the first-level contrast of parameter estimates masked by voxels activated in MT-on_TE=6.9_ and MT-off_TE=28(_ z > 3.1), as shown in figure S2. The MT-on data showed a similar temporal evolution of the hemodynamic response to the BOLD contrast found for MT-off.

### Data Analysis

MT-on and MT-off images were analysed using FSL 6.0.1 using the general linear model and FILM pre-whitening to find the activation in: (1) right visual cortex for left hemifield stimulation, (2) left visual cortex for right hemifield, and (3) their combination for both hemifields. The datasets were analysed using FSL FEAT^53^ to estimate the activation during the hemifield checkerboard task for all subjects and both conditions. The design matrix for the single-subject FEAT analysis modelled two explanatory variables (EV): (1) right hemifield trials vs. baseline, (2) left hemifield vs. baseline, each convolved with a hemodynamic response and constant term. Five contrasts were selected, [1 0] and [0 1] contrast for each EV to study the activation within each hemisphere, [1 1] contrast to model the common activation within the right and left hemifield. Finally, differential contrasts [1 -1] and [-1 1], were used to model specific activation within each hemisphere, excluding the common activation, herein referred to as (R-L) and (L-R). The last two contrasts detect activation unique to each hemifield-stimulus, and exclude non-specific responses to activation. The resultant z-score maps were combined to give the total specific activation maps.

The first-level analyses of all the subjects were combined into group-level analyses using fixed-effects, modelling the mean-difference across subjects, and mixed-effects (FLAME-1), allowing explicit modelling of both within- and between-subject variance components.^27^

To make inferences at the group level, a one-sided t-test was performed to assess the difference across methods. The one-sample t-test was conducted by concatenating parameter estimates from each contrast at the single-subject level. The z-(Gaussianised t/F) statistic images at the group-level were thresholded non-parametrically using clusters determined by Z>3.1 and a (corrected) cluster significance threshold of p=0.01.

## Supporting information

Suppementary Information

## Acknowledgements

The authors would like to thank Peter Hagoort, Laurentius Hüber, Peter Koopmans and Benedikt Poser for critical comments on this manuscript; Benedikt Ehinger and Floris de Lange for advice on stimulus design; and Christian Beckmann for advice on statistical analysis.

