## Supplementary material for "Arterial blood contrast (ABC) enabled by magnetization transfer (MT): a novel MRI technique for enhancing the measurement of brain activation changes": Suppementary Information

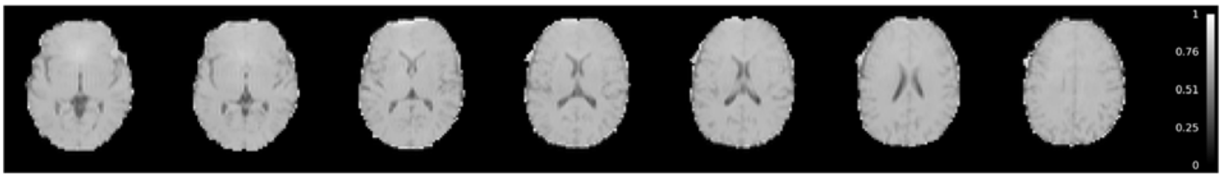


*Figure S1: Magnetization transfer ratio for a representative subject. The suppression factors are: grey matter / white matter (~60%, ~70 ); CSF suppression (~12-15%). Some residual effects of static field inhomogeneity are visible in frontal-temporal regions caused by the relatively long duration of the MT-pulses used in the present study.*

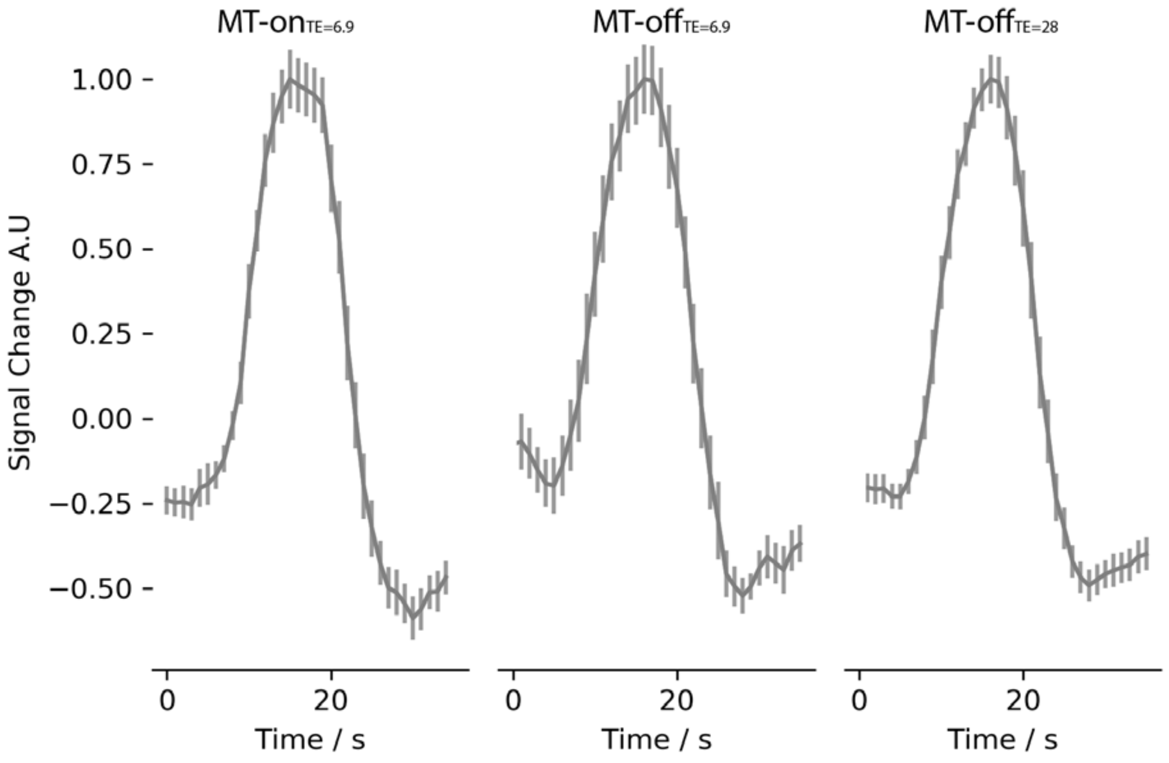


*Figure S2: From left to right, the hemodynamic response for MT-on$_{TE=6.9}$, MT-off$_{TE=6.9}$, and MT-off$_{TE=28}$. The hemodynamic response is generated within the visual cortex from the right and left hemifield, across 16 subjects using the ROI from the group-level z-score activation maps of BOLD contrast, thresholded at Z > 3.1. The results across subjects are highly reproducible (c.f. Figure S2). There is an indication that the hemodynamic response for MT-on has a broader top than MT-off.*

### Model of functional signal change as a function of TE

Following the analysis given in[1] we can ascribe the total signal coming from grey matter voxel to four compartments:

$$S(TE) = \sum_{i=1}^{i=4} x_i M_i exp(-R_{2i}^{*} TE)$$

Where the *i* correspond to: arterial, capillary, venous, and tissue compartments, the $x_i$ to the weighting factor, the $M_i$ to the magnetisation, and the $R_{2i}^{*}$ to the effective transverse relaxation rate of each compartment. Whereby

$$M_i = \frac{1 - exp(-R_{1i} TR)}{1 - \cos\alpha \, exp(-R_{1i} TR)} \sin\alpha$$

Where $R_1$ is the longitudinal relaxation rate and α the excitation angle, and

$$x_i = v_i C_i (1 - MTR)$$

$V_i$ is the volume fraction, and $C_i$ the fractional water content, and MTR the magnetisation transfer ratio. For simplicity we have introduced the MTR as a scaling factor rather than using the full Bloch equations.

The signal change upon activation will be given by

$$\Delta S(TE) = \sum_{i=1}^{i=4} x_{iact} M_{iact} exp(-R_{2iact}^{*} TE) - \sum_{i=1}^{i=4} x_{irest} M_{irest} exp(-R_{2irest}^{*} TE)$$

By taking values from the literature, we can plot the signal change as a function of TE for the situation both with and without MT.

The parameters that are independent of activation state are: $R_1$, $C_i$, and MTR. Whereby we have used: $T_{1blood}$=1650 ms;[2] $T_{1tissue}$=1600 ms;[3] $C_{blood}$=0.87; $C_{tissue}$=0.89; [1] $MTR_{blood}$=0.07; (scaled from[4]); $MTR_{tissue}$=0.4

Parameters dependent upon activation are: $v_i$, $R_{2i}^{*}$

Volume Fractions ($v_i$). We take the values given in[5] and assume a 30% increase upon activation.

| | Rest | Activation (30% increase) |
|---|---|---|
| Arteries | 0.0116 | 0.0203 |
| Capillaries | 0.0181 | 0.0224 |
| Veins | 0.0253 | 0.0288 |
| Tissue | 0.945 | 0.9285 |

Effective transverse relaxation rates ($R_{2i}^{*}$, Hz)

| | Rest | Activation |
|---|---|---|
| Arteries | 17.2 | 17.2 |
| Capillaries | 30.8 | 21 |
| Veins | 45 | 31 |
| Tissue | 15.2 | 14.8 |

We base the tissue T2* value for grey matter at rest as 66 ms given in,[6] and the change in T2* for parenchyma at 3T for visual stimulation as -0.38 Hz.[7] The R2* for arterial blood is taken from.[8] The venous R2* at rest is taken from[8] and the difference between rest and activation from.[9] The capillary values are based on Y=77 at rest and Y=91 at activation and calculated using (a*=16.75Hz, b*=37.6 Hz, c*=103) taken from.[8]

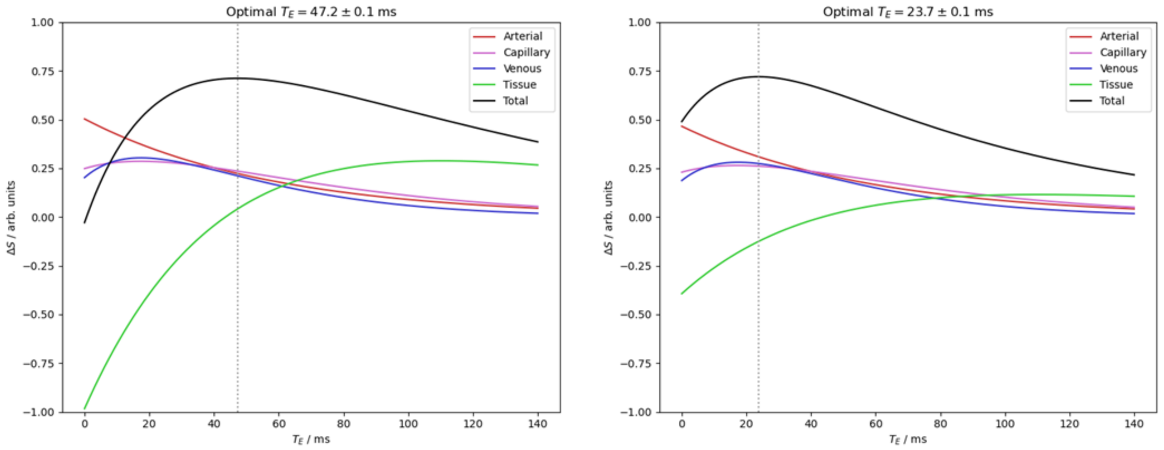


*Figure S3. The BOLD signal change is plotted as a function of TE for all four compartments, and the total BOLD signal change for MT-off (left) and MT-on (right). Arbitrary signal units are used, but the scales are the same for both figures. The optimal TE denotes that at which the total signal change is maximal. The tissue contribution to the total BOLD signal change is maximally negative at zero TE and increases thereafter. Reducing the tissue signal will increase the total signal change measured and reduces the TE at which the maximum signal change is recorded. The cross-over point where the total signal for MT-off becomes equal to MT-on is at TE=34.8 ms.*
